# Nanopore adaptive sampling: a tool for enrichment of low abundance species in metagenomic samples

**DOI:** 10.1101/2021.05.07.443191

**Authors:** Samuel Martin, Darren Heavens, Yuxuan Lan, Samuel Horsfield, Matthew D Clark, Richard M Leggett

## Abstract

**Background:** Adaptive sampling is a method of software-controlled enrichment unique to nanopore sequencing platforms recently implemented in Oxford Nanopore’s own control software. By examining the first few hundred bases of a DNA molecule as it passes through a pore, software can determine if the molecule is sufficiently interesting to sequence in its entirety. If not, the molecule is ejected from the pore by reversing the voltage across it, freeing the pore for a new molecule. User supplied sequences define the targets to be sequenced or ejected. Here we explore the potential of using adaptive sampling for enrichment of rarer species within metagenomic samples.

**Results:** We created a synthetic mock community consisting of seven bacterial species at different proportions ranging from 1.2% to 47% and used this as the basis for a series of enrichment and depletion experiments. To investigate the effect of DNA length on adaptive sampling efficiency, we created sequencing libraries with mean read lengths of 1.7 kbp, 4.7 kbp, 10.6 kbp, and 12.8 kbp and enriched or depleted for individual and multiple species over a series of sequencing runs. Across all experiments enrichment ranged from 1.67-fold for the most abundant species with the shortest read length to 13.87-fold for the least abundant species with the longest read length. Factoring in the reduction to sequence output associated with repeatedly rejecting molecules reduces the calculated efficiency of this enrichment to between 0.96-fold and 4.93-fold. We note that reducing ejections due to false negatives (approximately 36%) would significantly increase efficiency. We used the relationship between abundance, molecule length and enrichment factor to produce a mathematical model of enrichment based on molecule length and relative abundance, whose predictions correlated strongly with experimental data. A web application is provided to allow researchers to explore model predictions in advance of performing their own experiments.

**Conclusions:** Our data clearly demonstrates the benefit for enriching low abundant species in adaptive sampling metagenomic experiments, especially with longer molecules, and our mathematical model can be used to determine whether a given experimental DNA sample is suitable for adaptive sampling. Notably, repeated voltage reversals have no effect on pore stability.

## 1. Background

Whole genome shotgun sequencing of metagenomic samples has become a popular tool for understanding communities of mixed species. In particular, the ability to assemble individual species, or gene clusters such as antibiotic resistance genes, has the potential to shed new light on function, or to enable generation of reference sequences for unculturable organisms. With the increasing use of long read technologies, either on their own, or combined in hybrid approaches with short-read technologies, metagenome assembled genome (MAG) contiguity and accuracy metrics have improved still further^[1]^. Such approaches have been applied widely including in the assembly of pathogen genomes from clinical samples^[2]^, bacterial genomes and gene clusters from the human gut^[3]^, the rumen microbiome of cattle^[4]^, and a project which assembled tens of thousands of MAGs by re-analysing over 10,000 previously collected metagenomes^[5]^. Nevertheless, despite these successes, some doubts remain about the reliability of MAG approaches when faced with complex populations^[6]^.

Metagenomic samples are composed of a range of different species at varying levels of abundance. In nature, abundance often follows a power law^[7]^ and this can mean that sequencing of metagenomic samples produces data that results in deep coverage of some species with low or partial coverage of others. For rarer species, this is likely to result in much poorer assemblies and a reduction in the ability to distinguish between strains or related species. Effective enrichment strategies to maximise the sequence outputs of the rare species would address this weakness, and biodiversity blindspot.

Reducing host DNA is an important consideration in diagnostic applications, especially in clinical settings. A number of approaches are available as commercial kits, or detailed in published work including differential lysis, saponin-based depletion^[8]^, osmotic lysis^[9]^ or by enriching microbial DNA^[32]^. However, these approaches are not universally applicable, and require sample-specific adaptation often with many additional steps.

Hybridisation has been used effectively to both deplete and enrich samples prior to sequencing. For prokaryotes there are a number of commercially available ribosomal RNA depletion kits which claim to reduce the rRNA levels for transcriptomic studies from around 95% to less than 20%, but this still equates to a considerable amount of off target sequencing. Within eukaryotes as little as 2% of the genome can represent gene sequences coding for proteins. Exonic probe sets designed to target sequences of a few hundred base pairs have been widely used for mammals and plants and this strategy has been further optimised for molecules of a few thousand base pairs to determine gene structures when using approaches such as RenSeq^[10,11]^ when individual genes or families of genes are targeted. A common feature of these methods is an extended and often complicated library construction protocol which involves multiple PCR amplification steps that limits the length of DNA that can be interrogated. This can result in amplification biases in the output data and they require highly stringent hybridisation conditions coupled with accurate probe design to be effective.

In the same way that RNA sequencing can be compromised by highly expressed genes, metagenomic DNA samples can be compromised by highly abundant species. By adapting duplex specific nuclease based methods widely used for normalising cDNA libraries it is possible to normalise metagenomic samples. By denaturing and then slowly reannealing DNA molecules the likelihood is that the DNA from more abundant species are more likely to reanneal and become a substrate for the endonuclease^[12]^. However, problems can arise with non-specific annealing and with common regions such as the RNA operon which can share a high degree of similarity.

An additional development came with Clone Adapted Template Capture Hybridisation (CATCH-Seq) which was developed to resolve target regions of interest and circumnavigate the need to design specific probes. Using a bacterial artificial chromosome (BAC) known to contain regions of interest, generic probes are generated from the BAC and then used to pull down fragments spanning 60 kbp (single BAC) up to several hundred kbp (multiple BACs) to target difficult to sequence regions and help identify structural variation. Later protocols such as HLS-CATCH^[13]^ and nCATs^[14]^ use Cas-9 nuclease with guide RNAs to target DNA molecules up to several million base pairs in length.

An alternative to lab-based depletion or enrichment approaches is promised by Oxford Nanopore Technologies’ (ONT) adaptive sampling concept (sometimes called selective sequencing) which represents a form of software controlled enrichment. A programming interface known as “ReadUntil” enables control over individual pores and provides a mechanism for software to request ejection of the molecule currently being sequenced in a given pore. Thus the first few hundred bases of a molecule can be examined and a decision made if the molecule is ‘on target’. ‘Off target’ molecules are ejected by reversing the current across the pore, freeing the pore to capture a new molecule. In order for this to be effective, this must happen within a short time, so that the molecule can be ejected from the pore before most of it is sequenced. The longer the time taken for decision making and rejection, the lower the potential levels of enrichment possible.

Initially, ONT provided the ReadUntil programming interface and left it to third party developers to work out how to interrogate the raw pore signal to determine if a molecule was on-target. The first published implementation utilised a signal comparison algorithm known as Dynamic Time Warping (DTW) to compare the signal from the pore with pre-computed signals for sequences of interest^[15]^ (DTW also used in DySS^[16]^). However, this approach was computationally expensive, particularly for anything but relatively short reference sequences. Practical use was therefore limited due to the time required to make a decision when using larger reference databases. An alternative signal-based approach was provided by UNCALLED, which converted stretches of raw signal into k-mers and used higher probability k-mers as a query for a Ferragina-Manzini (FM) index search against a target database^[17]^. While more efficient than the DTW approach, it still required significant computational resources. RUBRIC^[18]^ abandoned signal-based comparison in favour of basecalling short (~150bp) portions of the start of reads and aligning to reference sequences using LAST^[19]^. However, this demonstrated limited enrichment and still required significant computational resources. More recently, ONT’s provision of real-time GPU-based basecalling on GridION devices enabled the development of Readfish^[20]^, which basecalls the first ~180 bases of sequence and aligns to references with minimap2^[21]^ in order to make a decision on accepting or rejecting a molecule. These solutions still required third party software in addition to ONT’s own control software. From the November 2020 release of the GridION control software, adaptive sampling was built in as a user-selectable option, which has opened it up to much wider adoption. The software requires a user to upload a file of reference sequences and the system can be set to either deplete or enrich for these.

Adaptive sampling offers a potential solution to enrich for species of interest in metagenomic samples. It requires a simple library construction method and samples can be processed within an hour without the need for amplification. However, a challenge for microbiome research is the difficulty of extracting high molecular weight DNA from metagenomic samples. Their unknown nature and the likely presence of both Gram positive, Gram negative and fungal species have led to the development of protocols such as the three peak extraction protocol where samples undergo three different methods involving either enzymatic, chemical or physical disruption to try and preserve DNA molecule length and ensure that the DNA faithfully represents all the species present in the sample^[22]^. This has shown that DNA molecules >20 kbp can be achieved, but for many scientists analysing microbiomes, bead beating is a necessity for DNA extraction due in part to the limitation of samples, the inability to effectively culture everything present and, in some cases, the need for rapid diagnostic results. This approach can be completed inside 20 minutes but typically produces DNA molecules <10 Kbp in length.

We wanted to investigate the effect of DNA molecule length on the efficiency and efficacy of adaptive sampling to determine its usefulness for both MAG and diagnostic applications. Here we present a mathematical model which can predict the enrichment levels possible in a metagenomic community given a known relative abundance and read length distribution. Using a synthetic mock community, we demonstrate that the predictions of the model correlate well with observed behaviour and quantify the negative effect on flow cell yields caused by employing adaptive sampling.

## 2. Results

### 2.1. A mathematical model of enrichment potential for metagenomic samples

A number of factors will affect the potential enrichment achievable through adaptive sampling. Here we derive a model to predict the theoretical enrichment of a set of taxa within a metagenomic community, based on the community composition and average read length. In the sections that follow, we show how this compares with real results achieved using the GridION. We begin with the assumptions (sequencing speed is 420 bases/s, the time taken to capture strands is 0.5 s and the response time once a strand is captured is 1.0 s) given in the worked example in the nanopore adaptive sampling information sheet^[23]^.

We can consider two alternative measures of enrichment: enrichment by yield and enrichment by composition. We define enrichment by yield as the ratio of the yields (per unit time) of target sequence (species) with and without adaptive sampling. This measure is likely to be the main consideration for researchers wishing to target particular sequences - if the overall target yield is less during targeted sequencing, then a better strategy would be to perform deeper untargeted sequencing and bioinformatically filter for the sequences of interest. One key factor that affects a nanopore sequencing run’s yield is the number of active pores. As the quality of pores before sequencing varies by flowcell, it is difficult to predict the yield of an experiment and compare adaptive sampling experiments between flowcells. Furthermore the use of protein pores is known to degrade them over time, possibly from the electric potential^[24]^, or from pore blockage^[17]^, thus repeated potential flipping from adaptive sampling could further decrease active pores and yield.

Enrichment by composition is the ratio of the relative abundance of target sequence (species) with and without adaptive sampling. This shows us how much the abundance of a given species in a metagenomic sample can be changed simply by employing adaptive sampling. This measure is not affected by yield, so we are able to use it to compare different experiments using different flowcells. By estimating the composition of target sequences in the community, it is then possible to estimate the target yield for a particular experiment design (assuming all flow cells being equally productive). Below, we develop a model to predict enrichment by composition.

Let *X* be the set of taxa present in a sample, and for a taxon *x* ∈ *X* define *x*_ab_ to be the abundance of *x* in terms of bases sequenced. This can be calculated by sequencing the sample without adaptive sampling, and calculating the sequence length of all reads that belong to the taxon *x*. Then the abundance of *x* can be given as this length divided by the total sequence length of all reads in the sample. For an experiment in which we enrich for *x*, let *x*_ob_ be the abundance of *x* observed in this experiment, calculated as before. Then we say that the enrichment factor for *x* (or simply, the enrichment of *x*) denoted *e_x_*, is given by

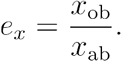

From this, it’s clear that the enrichment must be less than 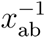, since *x*_ob_ ≤ 1 (for example, a taxon at 50% abundance cannot have an enrichment factor greater than 2). However, this fails to take into account the fact that in order to determine whether a read should be accepted or rejected, it has to be sequenced to some extent, and so we propose that the maximum achievable enrichment is in fact lower than this.

For more generality, we will partition *X* into the taxa to enrich (also called the target taxa), denoted *X*_e_, and those not to enrich (which by definition, will be depleted), denoted *X*_d_ = *X* \ *X*_e_. Following the ONT info sheet^[23]^, we estimate enrichment by the proportion of the total sequencing time that is spent sequencing the target taxa. We assume a constant sequencing rate throughout, denoted by *S* and in the units of number of bases sequenced per second. Let *T* be the proportion of sequencing time spent sequencing reads belonging to *X*_e_ without adaptive sampling, and *T*_e_ the proportion of time spent sequencing reads belonging to *X*_e_ with adaptive sampling. Then, since the sequencing rate is assumed constant, we can estimate the enrichment of *X*_e_ as

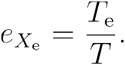

To determine the values *T*_e_ and *T*, we will fix the following quantities. Let *D* be the time taken between a molecule entering a pore and a decision being made on whether it should be accepted or rejected. Let *R* be the average read size, and let *C* be the time taken for a pore to capture a new molecule after sequencing a molecule. First we give an estimate for *T*. Denote by *y* the sum of abundances in *X*_e_, that is

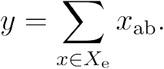

Each molecule takes, on average, *R/S* seconds to pass through the pore, and then a further *C* seconds until a new molecule is captured. The proportion of molecules that we want to enrich (i.e. to not eject from the pore) is *y*, so we have

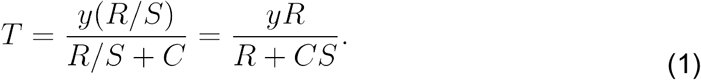

For *T*_e_, we make the following observation. For molecules belonging to *X*_e_ we still spend *R*/*S* seconds sequencing each molecule. For molecules belonging to *X*_d_ however, we spend *D* seconds sequencing each molecule. Thus the total sequencing time is given by *y*(*R*/*S*) + (1 – *y*)*D* + *C*, and so

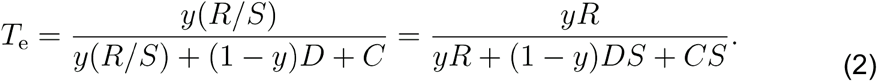

Taking the quotient of (1) and (2) gives us the formula for enrichment

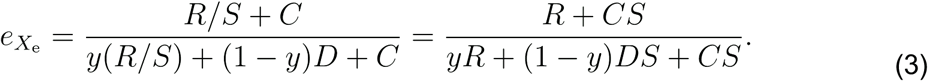

This formula gives us the enrichment of the whole set *X*_e_, but what if we want to determine the enrichment for a single taxon in *X*_e_? It’s an interesting feature of this model that it predicts enrichment to be the same for each taxon in *X*_e_. To see this, note that if we were to determine the enrichment of a single taxon *x* ∈ *X*_e_, then in equations (1) and (2), we would replace *y* in the numerator with *x*_ab_, whilst the denominators remain the same. But then, in taking the quotient in equation (3) these terms cancel.

If we wish to enrich for a taxon *x* ∈ *X* (so that *X*_e_ = {*x*}), then we have that *y* = *x*_ab_ and equation (3) becomes

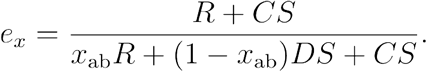

We can rewrite the denominator as *x*_ab_(*R* + *CS*) + (1 – *x*_ab_) (*DS* + *CS*). Since 0 < *x*_ab_ < 1 and *C*, *D*, *S* > 0 we have that *x*_ab_(*R* + *CS*) + (1 – *x*_ab_)(*DS* + *CS*) > *x*_ab_(*R* + *CS*), and so

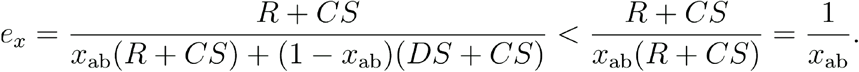

Thus our model predicts that enrichment of a single species will be lower than 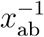, as discussed at the beginning of this section.

We created a Shiny web application which allows researchers to explore the potential enrichment that may be possible for their experiments. The app can be found at https://sr-martin.shinyapps.io/model_app/ and allows the user to adjust parameters such as the average read length to explore the effect on enrichment potential for species of varying abundance.

### 2.2. Starting relative abundance and molecule length determine the level of enrichment

We created a bacterial mock community consisting of seven species ranging in abundance from just over 1% to around 47% (Table 1) as determined during control sequencing runs. In order to observe the effect of molecule length on enrichment, we performed a series of experiments with different library preparations, each producing a different read length from the same input material (Table 2). For simplicity, we refer to the library by the mean read length generated during control runs; however, this value alone is insufficient to capture the sometimes complex read length distribution of the library (Figures 1a,b). For each library, we performed a control run in which we sequenced for approximately 1 hour (enough for at least 17,086 reads, and averaging 70,420). We then enabled Adaptive Sampling and enriched for each bacterial genome, one by one, starting with the most abundant species and ending with the least abundant. For the library with a mean of 10.6kbp, we performed an additional “Low to High” run, in which the bacteria were enriched in reverse order, lowest abundance first. For both of these runs, we maintained half the pores as control pores throughout; for all other runs, we did not maintain control pores after the initial control run.

**Table 1:**
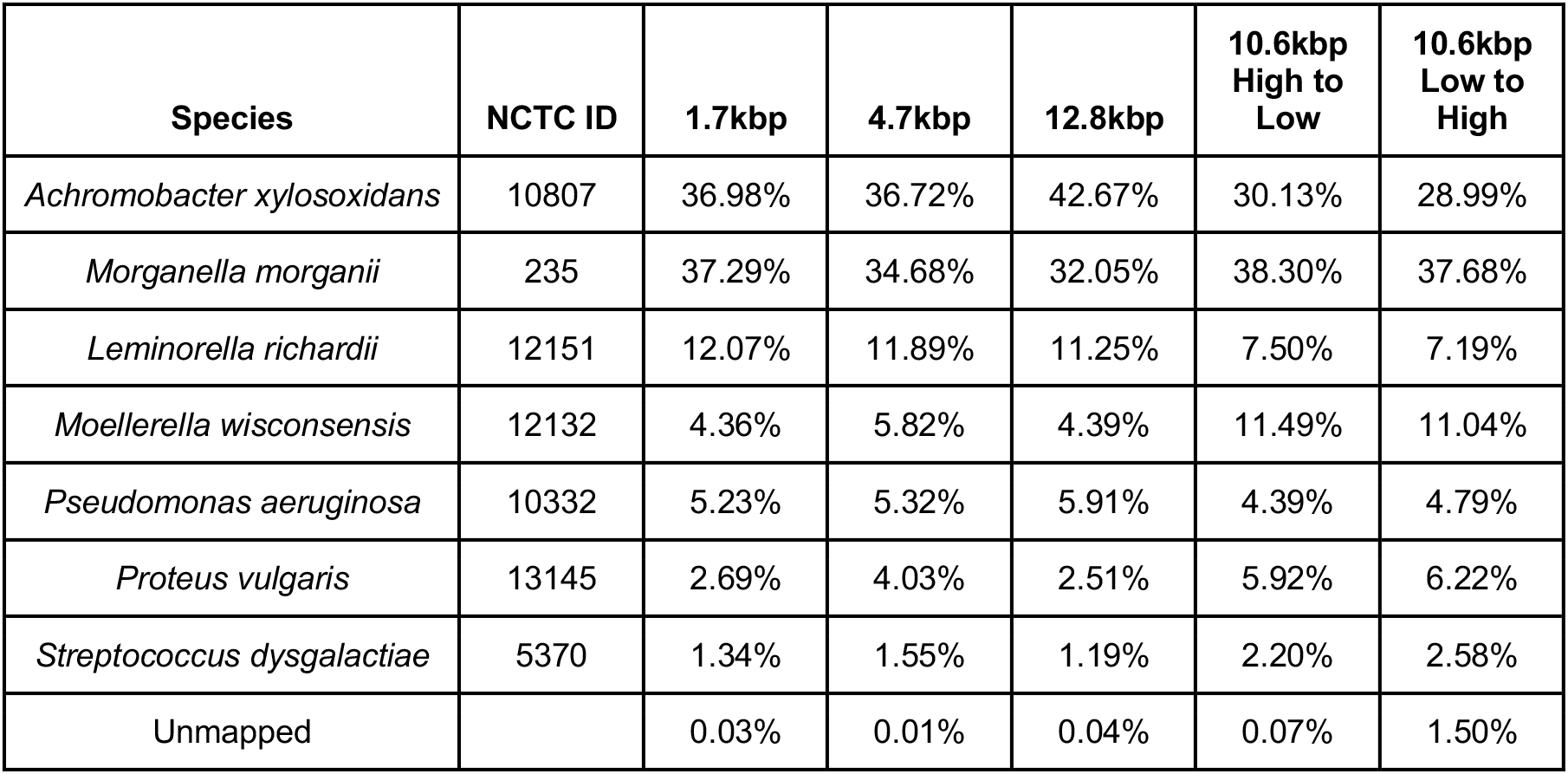
Relative abundance of the 7 bacteria used in the mock community, as determined from control runs. All were selected from the National Collection of Type Cultures and strain IDs are given. Percentages represent the percentage of sequenced bp aligned to reference genomes. Read counts give similar percentages and can be found in Supplementary Table 1.

**Table 2:**
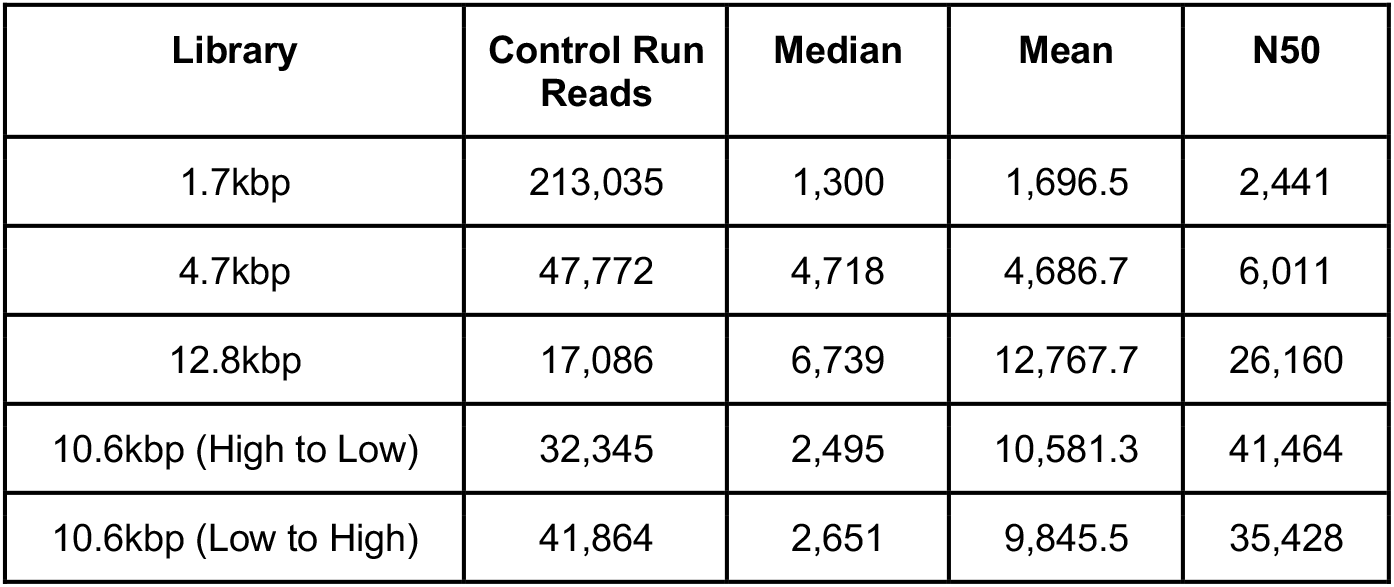
Read statistics for control runs for each library.

**Figure 1.**
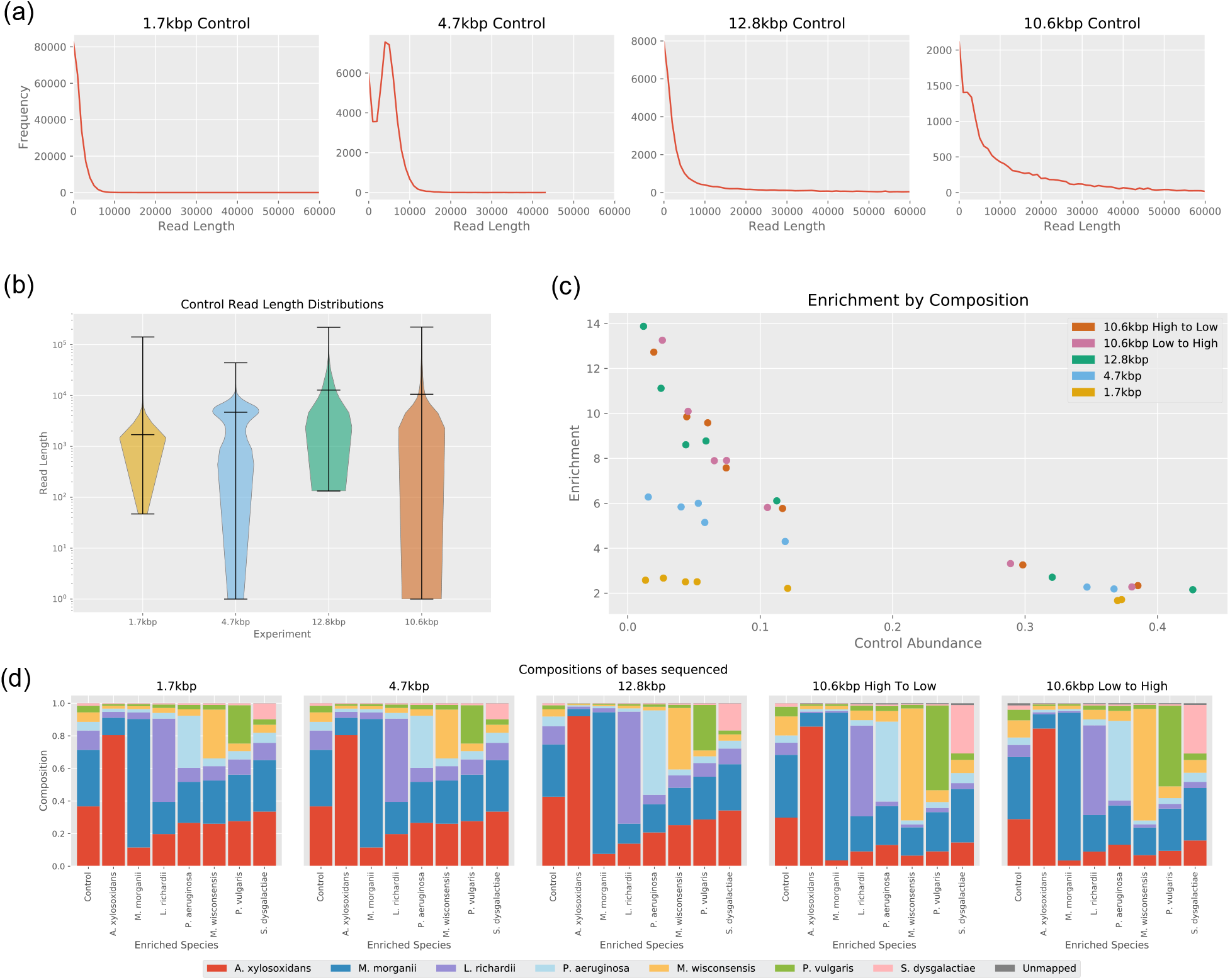
(a) Read length distributions from control runs. Reads were binned by length into bins of size 1000bp. Distribution for 10.6kbp library taken from control run in High-to-Low experiment. (b) Violin plots (log scale) of read length distributions from control runs. Distribution for 10.6kbp library taken from control run in High-to-Low experiment. Extrema and means shown in black. (c) Enrichment factor against relative abundance. Each point represents a species, with the position on the x axis indicating the original relative abundance of the species and the position on the y axis indicating the enrichment factor obtained. (d) Community composition for each enrichment target during the runs.

We calculated the enrichment by composition by dividing the relative abundance of a species with enrichment by the relative abundance without enrichment. As predicted by the model, the enrichment factor was higher for less abundant species, and for longer read lengths (Figure 1c). Highest levels of enrichment were produced for *S. dysgalactiae*, with relative abundance changed from 1.19% to 16.52%. The effect on community composition can be seen in Figure 1d.

### 2.3. Enrichment by composition approaches the maximum predicted by the model

We compared the model predictions with the results of the mock community runs. For the 1.7kbp, 4.7kbp and 12.8kbp runs, we calculated enrichment by composition as the quotient of the abundance during enrichment and the abundance during the control run. For the 10.6kbp runs, enrichment by composition was calculated as the quotient of the abundance on enrichment channels (1-256) and abundance on control channels (257-512) for each species in the mock community. Following the ONT info sheet^[23]^ we used the fixed values *S* = 420bps (bases per second), *C* = 0.5s, and *D* = 1.0s. Figure 2a overlays results from the experimental runs with predictions from the model.

**Figure 2:**
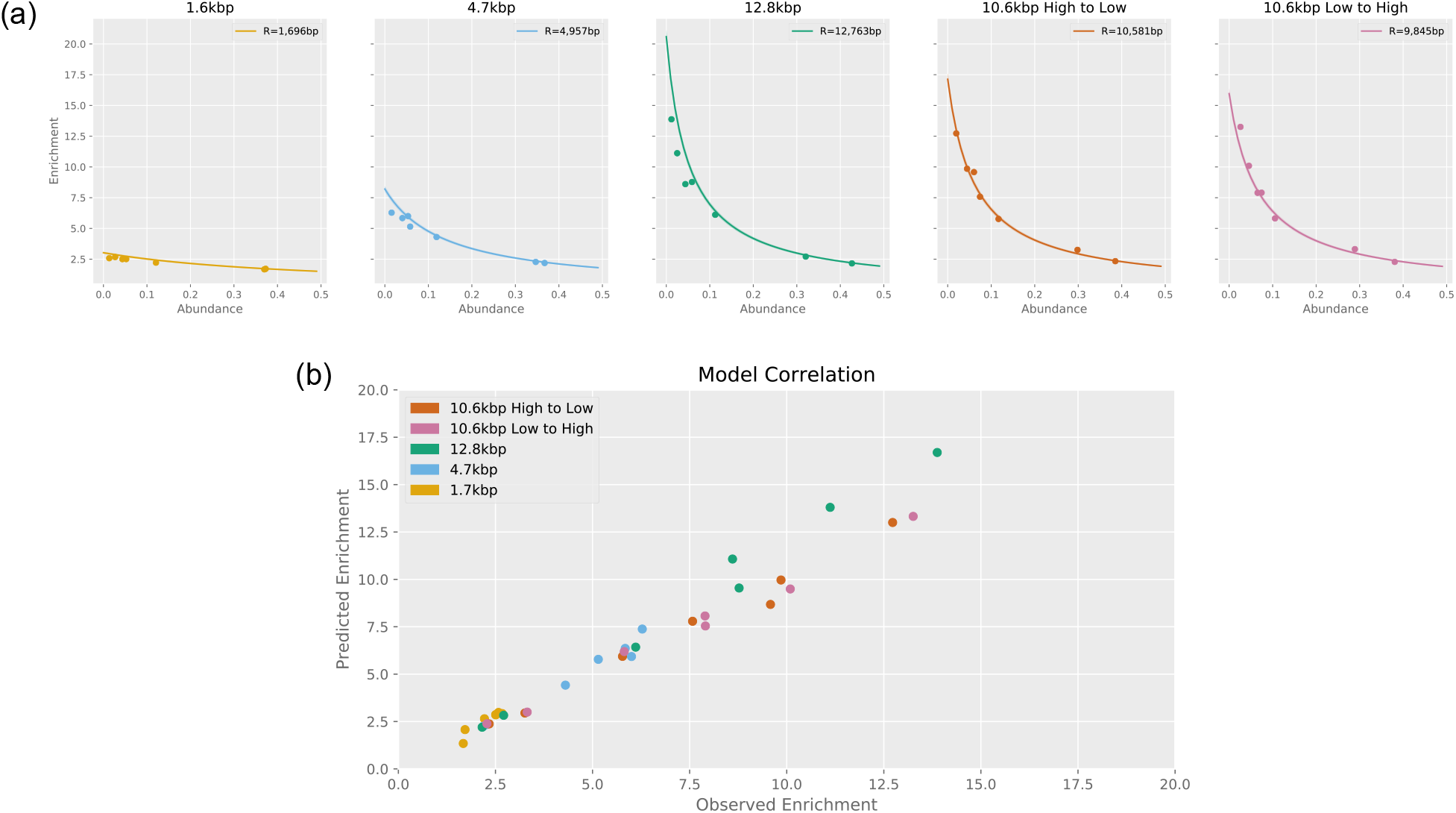
(a) Scatterplots of enrichment vs abundance. Curves show enrichment values predicted by the model for average read lengths. (b) Correlation between observed enrichment values and predicted enrichment (Pearson’s r of 0.9825).

We calculated the root-mean-square deviation of each data set from the values predicted by the corresponding model (Table 3). Our model predictions also correlated strongly with our observations (Pearson’s r = 0.9825) as can be seen in Figure 2b.

**Table 3:** Root-mean-square deviation of experiments from model.

| Run Name | Mean Read Length | Root-mean-square deviation |
| --- | --- | --- |
| 1.7kbp | 1,696 | 0.350 |
| 4.7kbp | 4,957 | 0.518 |
| 12.8kbp | 12,763 | 1.821 |
| 10.6kbp High to Low | 10,581 | 0.392 |
| 10.6kbp Low to High | 9,845 | 0.333 |

### 2.4. Enrichment by yield is significantly less than enrichment by composition

For each run, we also calculated enrichment by yield. For the 1.7kbp, 4.7kbp and 12.8kbp runs, we calculated enrichment by yield as the quotient of the yield per hour per active channel during enrichment and the yield per hour per active channel during the control run. For the 10.6kbp runs, enrichment by yield was calculated as the quotient of the yield per hour per active channel on enrichment channels (1-256) and yield per hour per active channel on control channels (257-512), for each species in the mock community. For the 1.7kbp run, yield of target sequences was slightly lower during adaptive sampling than during the control run (Figure 3a). Normalising the yield by the number of active channels during the first 30 minutes of each experiment confirms this (Figure 3c). For the 1.7kbp and 4.7kbp runs, we performed another control experiment after the adaptive sampling. Figure 3a,b,c indicate significantly reduced yield after adaptive sampling than the yield before adaptive sampling, particularly for the 1.7 kbp run.

**Figure 3:**
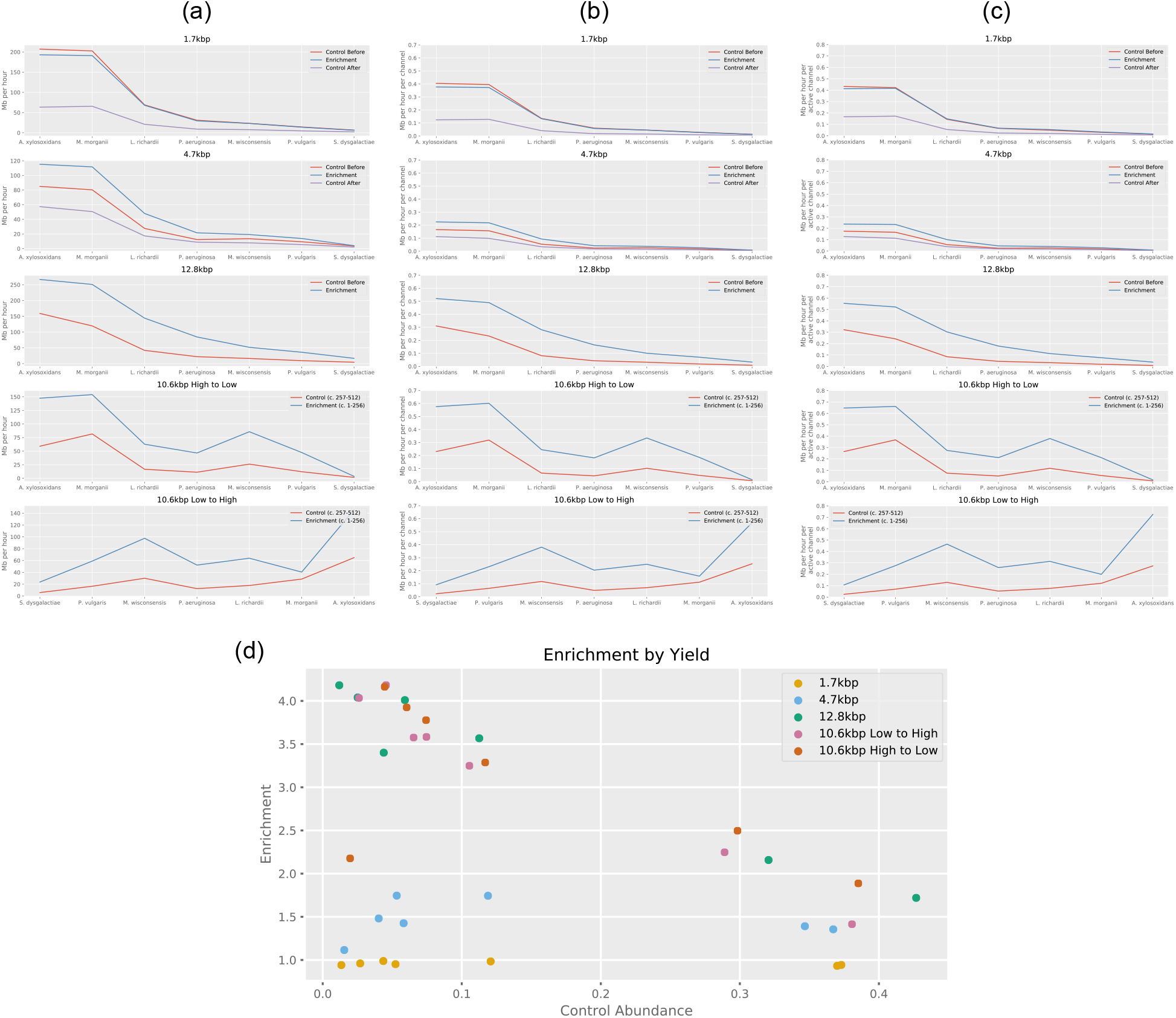
(a) Yield of target sequences in Mb per hour during adaptive sampling (blue), control before/during (red), and control after (purple). (b) Yields per hour for all runs, normalised by channels used. (c) Yield of target sequences in Mb per hour per active channel during adaptive sampling. (d) Enrichment by yield values. Each experiment, except for the 1.7kbp run, gave us increased yield when performing adaptive sampling.

Figure 3d summarises the levels of enrichment found for all bacteria in all runs. Highest enrichment of 4.93x was found for *P. aeruginosa* in the 10.6kbp Low to High run.

### 2.5. Reducing false negative identifications and associated pore ejections would significantly increase enrichment by yield

It’s apparent from Figure 2a that the observed enrichment for low abundance species during the 12.8kbp run was less than the model predicted. Figure 4a shows the distribution of read lengths for the control portion of this run.

**Figure 4:**
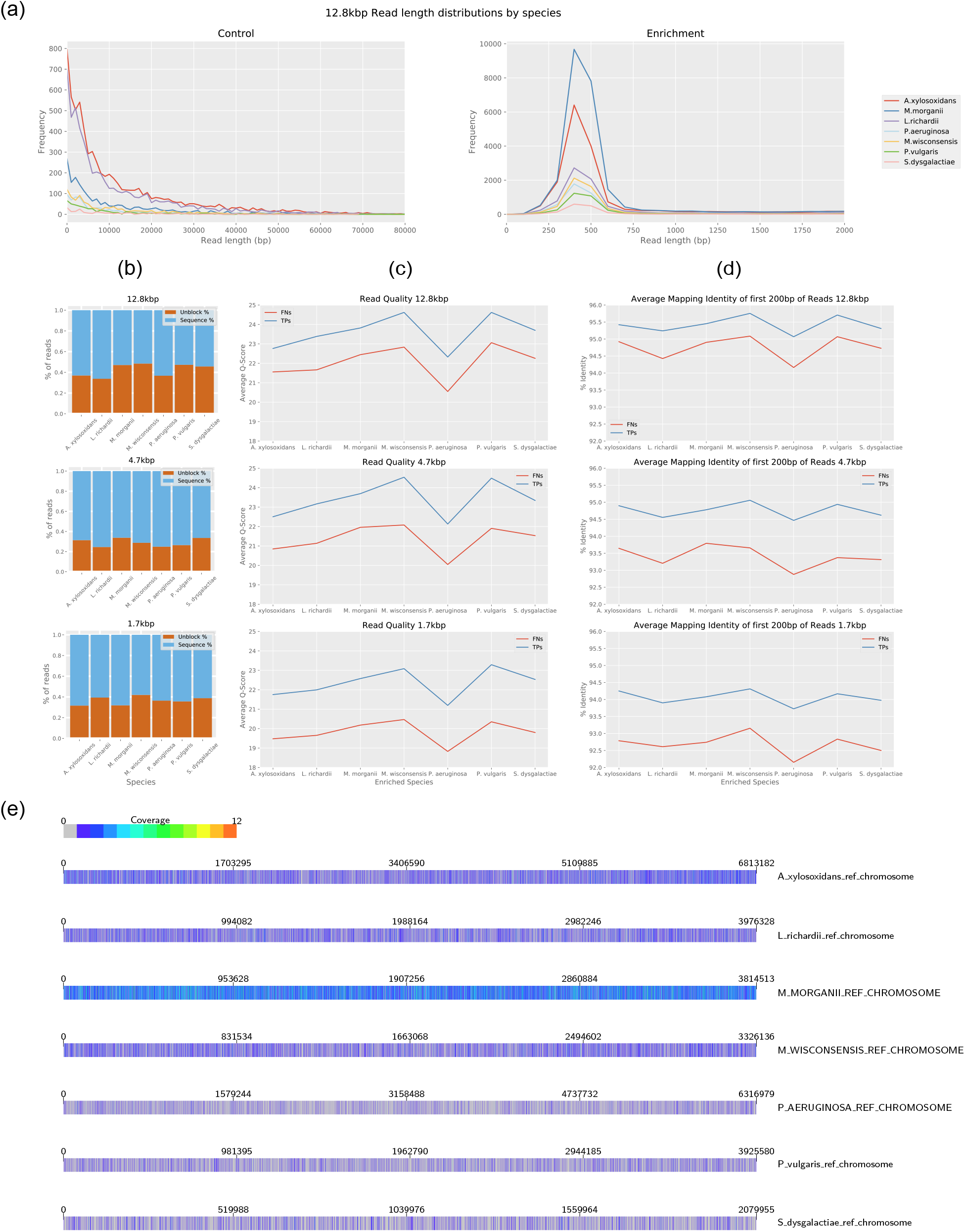
(a) Distribution of read lengths during control portion and enrichment portions of 12.8kbp run. Reads are split by species. (b) Proportion of target reads rejected during adaptive sampling. (c) Quality values of reads, split by species and TP/FN. (d) Average identity of mappings of first 200bp of reads against reference genomes. The mapping to the correct genome with the highest identity was used to calculate the averages. (e) Coverage of target genomes by false negative reads (i.e. reads there were incorrectly ejected from the pore during adaptive sampling) during 12.8kbp run. Image produced using the alignment visualisation software Alvis^[26]^.

During adaptive sampling we expect to see distributions similar to these for species that are being enriched, and a sharp peak around 500bp for all other species, which are depleted. However, we find that, when a species is being enriched, it also displays a peak around 500bp, suggesting that target molecules are being rejected (Figure 4a).

By parsing the logs provided by the GridION, we found that during adaptive sampling, approximately 36% (lowest 24.6%, highest 48.5%) of target molecules were being ejected from the pore. These are molecules that are misclassified as non-targets by the first fast mapping, but subsequently classified as targets by post-enrichment alignment (Figure 4b). We performed further analysis to determine why this was. We split the read sets first by their species classification, and then by the signal sent to the pore when they were sequenced. Thus each species had a set of sequenced reads (true positives) and ejected reads (false negatives). First, we calculated the average quality score of each read (Figure 4c). This shows that the average quality of TPs was higher than the average quality of FNs for each run. Next, we took the first 200bp from each read and used BLAST^[25]^ to map them against the reference genomes. By doing this, we are attempting to use just the sequence data that is available to MinKNOW when it makes a decision during sequencing on whether to sequence the molecule, or eject it from the pore. For each read we took the mapping of its first 200bp which had the highest identity and mapped to the correct genome, and used these to calculate the average identity (Figure 4d). We found that the TPs had a higher average identity than the FNs, although in this case the TPs for the 1.7kbp run had a lower identity than the FPs from the 12.8kbp run. To determine whether regions of low genome complexity can affect the FN rate, we mapped all FN reads to their true target genome. A heatmap showing the coverage of each genome by the FN reads is displayed in Figure 4e and shows no obvious clustering.

### 2.6. Use of adaptive sampling has an effect on active pores, but increases target yield and MAG assembly potential

Our experiments demonstrated continued enrichment over 8-9 hour sequencing periods, but we wanted to see how the repeated rejection of molecules affected the lifespan of the pores and if enrichment was still worthwhile over longer periods of time. We ran two new flowcells in which we enriched for a low abundant species (*S. dysgalactiae*, ~2.6%) and a high abundant species (*M. morganii*, ~37.5%). We also ran two flowcells in which we depleted *M. morganii* (~ 37.5% of total) and depleted *M. morganii, A. xylosoxidans* and *L. richardii* (together ~ 74.2% of total). With all four flowcells, half the channels were used as control channels in which no adaptive sampling took place. As previously, all four flowcells demonstrated increased yield of the target species, but a decreased total yield (Figure 5). The number of active channels was slightly higher for control channels than for enriched channels, but the difference was not large (Figure 6b). Hourly yield for target species was consistently higher for the first 24 hours with adaptive sampling (Figure 6c). However, yield of target species declined at a greater proportionate rate on the adaptive sampling channels (down 36% from hour 1 to hour 6) than the control channels (down 25% between hour 1 and hour 6). By 50 hours, hourly yield for adaptive sampling was similar to the control channels, but overall flow cell life was much declined by this point, in line with expectations for current nanopore flow cells. Time between target reads was reduced considerably in adaptive sampling channels over the control channels (Figure 6d).

**Figure 5:**
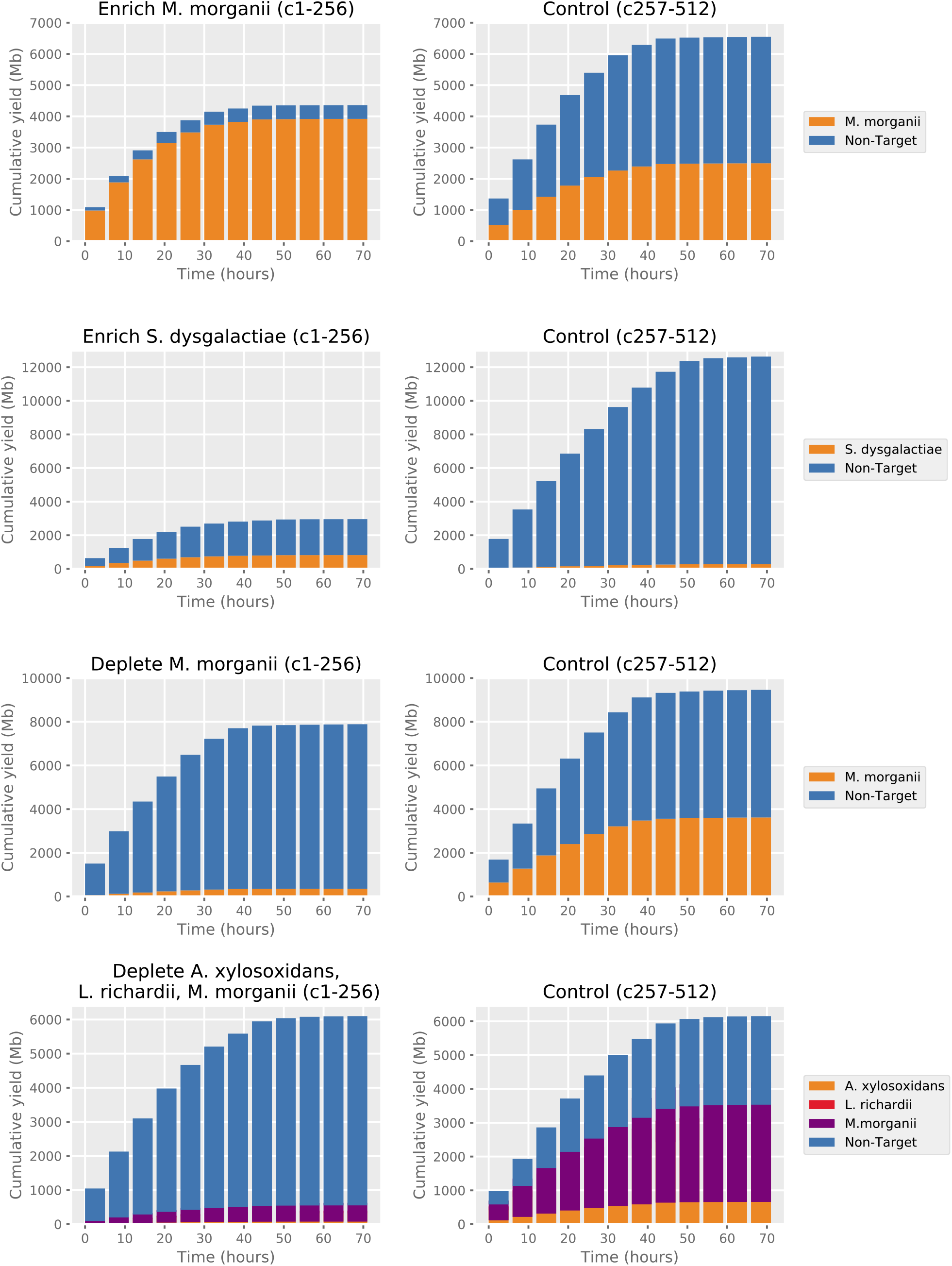
Cumulative yields split by experiment channels and control channels.

**Figure 6:**
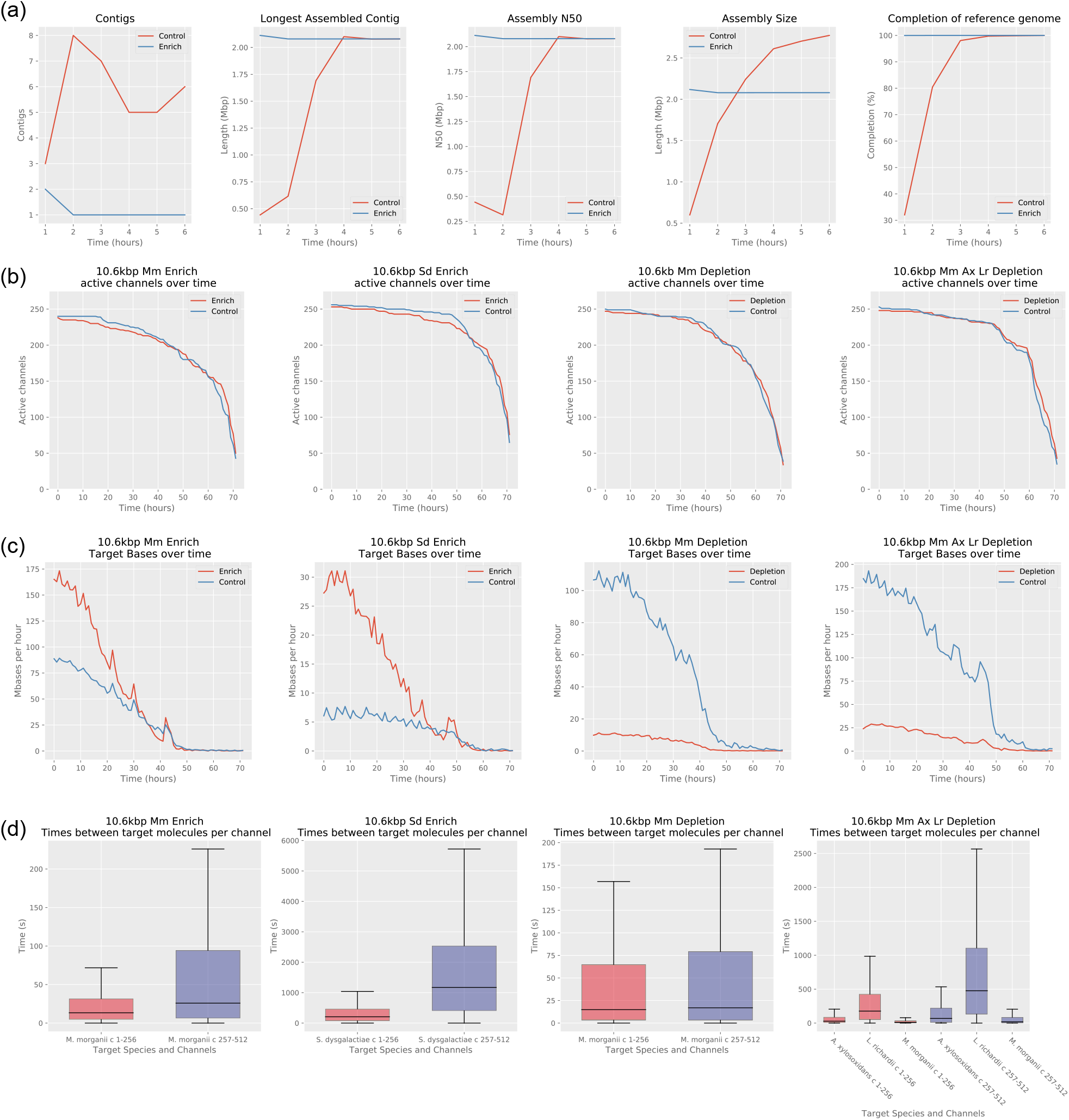
(a) *S. dysgalactiae* assembly statistics for enriched and control channels. (b) Plots showing how the number of active channels varies with time. (c) Hourly yields from enriched/depleted channels vs control channels. (d) Times between consecutive target molecules on individual channels, split by enrich/deplete (channels 1-256, red) and control (channels 257-512, blue).

Reasoning that one mechanism of pore loss is clogging by DNA that can’t be ejected^[17]^ we tested a nuclease flush during a sequencing run for a possible recoverative effect on pores used for adaptive sampling. We ran a new flowcell enriching for a single low abundant species (*S. dysgalactiae*, ~2.6%) for 6 hours, carried out a nuclease flush and then ran the flow cell for a further 6 hours. The flush appeared to result in an increased number of active channels for both the control and enriched portions of the flowcell (Figure 7a). The effect on hourly yield was less clear (Figure 7b).

**Figure 7:**
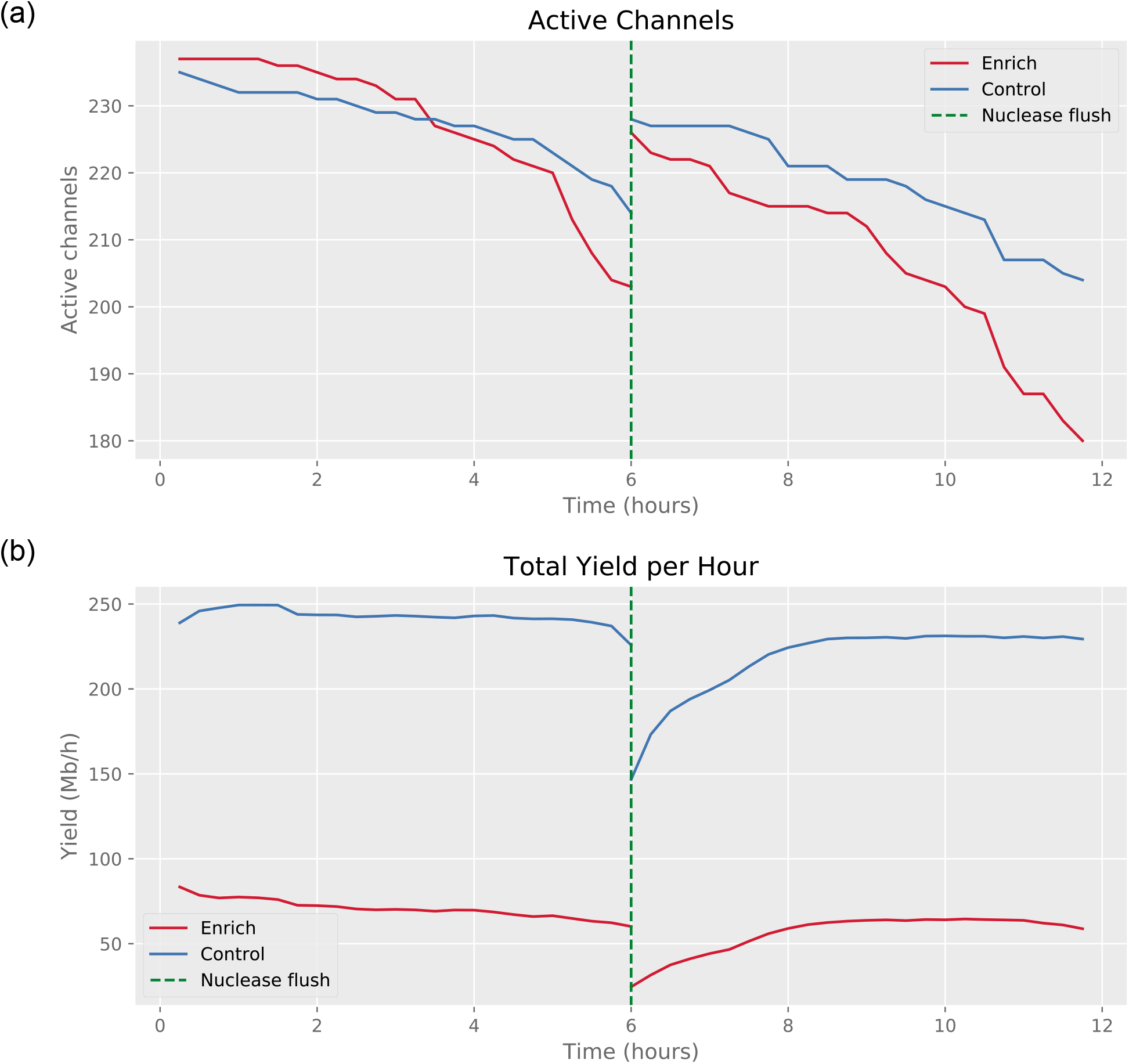
Effect of nuclease flush on active channels and yield.

In order to evaluate the effect of adaptive sampling on the potential for MAG assembly of a low abundance species, we took the reads available at 1 hour intervals from the control channels and the enriched channels. Reads mapping to the *S. dysgalactiae* reference were used as the input to the Flye assembler^[27]^. For the enriched channels, a single contig, high accuracy assembly was produced with the data available at 2 hours (Table 4, Figure 6a). Subsequently, we also performed an assembly of the enriched channel with reads available at 1.5 hours and this also produced a single contig assembly. For the control channels, after 6 hours, the *S. dysgalactiae* yield (32 Mbp) had not yet reached that produced by the enriched channels in 1.5 hours (42 Mbp), which was also reflected in much lower contiguity (6 contigs vs 1 contig).

**Table 4:**
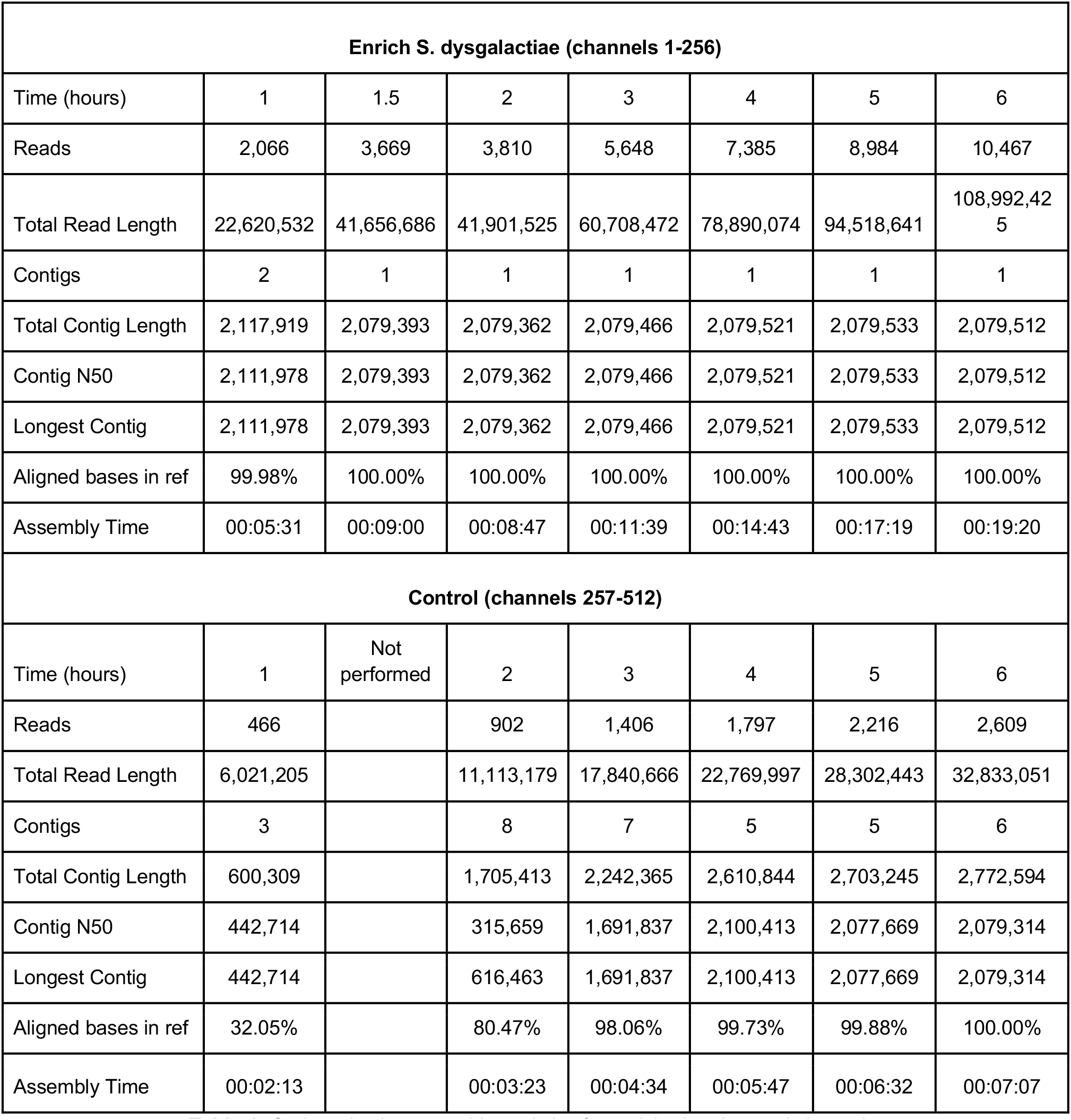
*S. dysgalactiae* assembly statistics for enriched and control channels.

## 3. Discussion

Previous studies have demonstrated the use of bespoke third party adaptive sampling software to enrich for sequences within an organism e.g. exons or loci of key variants. Here we test ONT’s own recent implementation of adaptive sampling in the GridION control software as a tool for enriching or depleting species in metagenomic samples. We describe a mathematical model that can predict enrichment potential for a species of given relative abundance and mean read length, and show that enrichment by composition in real experiments closely follows that predicted by the model (Pearson’s r of 0.9825). Enrichment by yield, the value that is of most practical benefit to researchers, lags behind enrichment by composition, but we show that with longer read lengths, we were able to enrich relatively low abundance (~2%) organisms by almost 5x. High quality single contig MAG assemblies of the same species were possible within 1.5 hrs using adaptive sequencing and around 6 hours without. Adaptive sequencing could be fed with MAG sequences (from existing short read assemblies), verifying true assemblies, splitting chimeric MAGs^[28]^ (using long nanopore reads as an orthogonal data type), or improving assemblies using long read scaffolding.

Two key factors affect the enrichment potential. Firstly, the initial abundance of the target species determines the theoretical maximum levels of enrichment. Secondly, the length of DNA molecules presented to the sequencing pores limits the efficiency of the process. For shorter molecules, the time taken to basecall, align, reject a molecule and capture an alternative becomes significant compared to the time taken to sequence a typical read without adaptive sampling. It is for this reason that adaptive sampling of our 1.7kbp mean library was not beneficial and possibly slightly detrimental to overall yield. Yet for longer reads, there are clear benefits to the adaptive sampling approach. Our data also indicate that there is still much potential to improve on current adaptive sampling implementations and to increase the enrichment by yield significantly. Our analysis showed that around 40% of on-target reads were falsely rejected. Where this value was highest, in the mean 10.6kbp run, we observed slightly lower enrichment by composition values than those predicted by our model, and believe these incorrect rejections to be an explanation for the difference. Reducing incorrect ejections could increase target yield significantly and thus result in much higher enrichment factors. ONT provide two implementations of their GPU basecalling algorithm - a faster, less accurate one and a slower, more accurate approach. The adaptive sampling basecalling is performed with the higher speed, lower accuracy algorithm. When hardware progression or algorithmic improvements enable the use of the more accurate basecalling algorithm, this will likely bring a reduction in false negative ejections. Additionally, improvements to the alignment and decision making approaches employed, as well as to the underlying ReadUntil API, will also bring improvements.

In evaluating the use of adaptive sampling in a particular metagenomic application, a prime consideration will be the ability to prepare DNA that is long enough to derive meaningful enrichment. ONT and a number of users have demonstrated megabase read lengths from genomic samples^[29,30]^, but it is not possible to imagine such read lengths in complex metagenomic samples due to the need to lyse different cell types, including some that are particularly troublesome to break open. A move away from mechanical lysis approaches such as bead beating towards newer enzymatic techniques will be key. The desire to target particular species with known cell wall characteristics - e.g. for assembly - may mean that harsher lysis approaches are unnecessary for that species. Even with bead beating, we have previously demonstrated DNA extractions from faeces (not the simplest of samples) can produce nanopore data with mean read lengths as high as 8.1kbp^[2]^, and in other experiments we have generated mean read lengths of up to 15kbp from soil metagenome samples (Heavens D, unpublished data). Our data indicates that significant enrichment is achievable at these lengths. If input material is not limited, then physical (bead or gel based) size selection can be used to increase the mean read length further. Even the 4.7kbp run presented here produced enrichment of 2x, which could mean experiments cost half as much money or take half as long to complete. This reduction in time could be particularly important in clinical applications or for environmental pathogen detection.

Previous studies have shown a faster decline in active sequencing channels for flowcells undergoing adaptive sequencing, which may be due to the act of rejecting molecules or that the likelihood of clogging is statistically related to the number of molecules captured by a pore^[17,20]^. Our own data shows a slight decline in active channels (Figure 6b), but not as much as seen previously. Nevertheless, like others we find that overall yield including non-target yield is reduced compared to control channels (Figure 5), particularly when enriching for lower abundance organisms. We find that a nuclease flush appears to have a restorative effect on active channel count, but the effect on yield was less clear. It’s possible that 6 hours was too soon to derive much benefit and others have suggested flushing every 24 h^[20]^.

Overall, our results show that adaptive sampling can increase target yield significantly in real terms, provided that molecules of a modest length are used. Given the strong effect of read (library molecule) length it is likely that ONT ligation based libraries would outperform rapid based ones from the same material, as the transposase would decrease library molecule size. The use of adaptive sampling provides us with the benefits of library-based enrichment, without complex protocols or the bias that these may introduce. This is a significant advantage to researchers who may not have access to specialised laboratory equipment. Furthermore, it maintains the advantages of nanopore sequencing e.g. speed, longer read lengths, detecting methylation and other epigenetic modifications using the raw nanopore signal, and the possibility of conducting experiments in-field.

We envisage several applications of adaptive sampling in the near future. One possibility is the targeting of molecules to close gaps in reference genomes. This could be achieved by enriching for molecules that align to sequences flanking gaps in the genome, and depleting everything else. Whilst we demonstrated over 4-fold enrichment in terms of yield, the potential read lengths for metagenomic applications are limited by the variety of DNA extraction methods required for the many cell types that may be present in the sample. Significantly higher average read lengths are possible for non-metagenomic samples, and so the potential for enrichment is greater.

Another possible application of adaptive sampling is the improvement of MAGs. In Section 2.6 we demonstrated the improved time-to-assembly of a known bacteria using adaptive sampling. In the future, we plan to develop a pipeline to assemble metagenomic reads *de novo* in real time during the experiment. Using adaptive sequencing, we could deplete molecules that cover regions that are already well assembled, or even enrich for reads with sequence at the ends of contigs and pointing into the unknown region, maximising the useful data to improve the assembly. Even rejected reads (~500bp) need not go to waste as these can be used for digital abundance measurements and for polishing assemblies. Existing software such as Readfish^[20]^ already enables the updating of target regions during a run, thereby allowing continual adjustment of targets to refine the assembly as the sequencing progresses. We believe this would lead to improved MAG quality, particularly for low abundance species.

## 4. Conclusions

Through ONT’s adaptive sampling software, we demonstrated enrichment in terms of both yield and composition, in a synthetic mock metagenomic community. We found that enrichment was higher for lower abundant species, and for libraries with a higher average molecule length, showing that extraction methods that can preserve molecule length are key to obtaining the highest enrichment. We developed a mathematical model to estimate the enrichment by composition that can be expected based on experimental factors, and showed that the model’s predictions correlated strongly with the observed data. We also observed that the occurrence of false negatives affects the achieved enrichment, but expect that improvements in hardware and software will minimise this in the future. By performing targeted enrichment on a low abundance species, we were able to significantly reduce the time taken to achieve a high-accuracy, single contig assembly, compared to non-targeted sequencing. Notably, we found that the repeated ejection of molecules from the pores had less effect on pore stability than has been previously reported. We conclude that adaptive sampling will prove to be a useful tool for many nanopore-based metagenomic studies.

## 5. Methods

### 5.1. Bacterial cell culture and DNA extraction

Seven bacterial strains were identified from the NCTC that had a fully assembled single chromosome genome, varying GC content and sizes with no plasmids. Full strain details and assemblies available at https://www.sanger.ac.uk/resources/downloads/bacteria/nctc/. Bacteria were grown overnight in 3ml of 2xYT in a 5ml tube in an Eppendorf Thermomixer C at 37°C shaking at 400rpm. Following the incubation the tubes were spun at max speed in an Eppendorf 5427R centrifuge for 5 minutes to pellet the cells and the supernatant discarded.

For the Gram positive bacteria cell pellets were resuspended in 160***μ***l of Qiagen P1 buffer, transferred to a 1.5ml tube and then 20***μ***l of 100mg/ ml lysozyme added, mixed and incubated for 30 minutes at 37°C shaking at 900rpm in an Eppendorf Thermomixer C. To this 20***μ***l of proteinase K was added and incubated for 30 minutes at 56°C shaking at 900rpm. The tube was then cooled on ice and 2***μ***l of RNase added and incubated at room temperature for 2 minutes.

To precipitate the DNA onto beads 150***μ***l of ATL was added followed by 15***μ***l of MagAttract Suspension G and 280***μ***l of MB buffer. This was incubated for 3 minutes at room temperature shaking at 1 400rpm. The beads were then pelleted on a magnetic particle concentrator (MPC), the supernatant discarded and the beads washed twice with 700***μ***l MW1 buffer and twice with PE buffer resuspending the beads on each occasion.

Two 700***μ***l water washes were then performed whilst the beads remained on the MPC incubating for 1 minute at a time. DNA was then eluted from the beads by mixing the beads for 3 minutes at room temperature shaking at 1 400rpm in 100***μ***l AE buffer.

For the Gram negative bacteria cell pellets were resuspended in 180***μ***l of ATL buffer, transferred to a 1.5ml tube and 20***μ***l of proteinase K was added and incubated for 30 minutes at 56°C shaking at 900rpm. The tube was then cooled on ice and 2***μ***l of RNase added and incubated at room temperature for 2 minutes.

To precipitate the DNA onto beads 150***μ***l of ATL was added followed by 15***μ***l of MagAttract Suspension G and 280***μ***l of MB buffer. This was incubated for 3 minutes at room temperature shaking at 1 400rpm. The beads were then pelleted on a MPC, the supernatant discarded and the beads washed twice with 700***μ***l MW1 buffer and twice with PE buffer resuspending the beads on each occasion.

Two 700***μ***l water washes were then performed whilst the beads remained on the MPC incubating for 1 minute at a time. DNA was then eluted from the beads by mixing the beads for 3 minutes at room temperature shaking at 1 400rpm in 100***μ***l AE buffer.

### 5.2. DNA QC

DNA concentration was determined using the Life Technologies Qubit broad range and high sensitivity assay kits. A 1 ***μ***l aliquot of DNA was combined with 198***μ***l of the appropriate buffer and 1 ***μ***l of dye in a 0.5ml qubit tube, vortexed and left at room temperature for 2 minutes. DNA concentration was then measured on a Qubit 3 fluorometer. If DNA concentration between the high sense and broad range assays differed by more than 10% then the extractions were repeated. DNA was then calculated by averaging the measurement from each assay.

To confirm molecule length extracted DNA was run on either the Agilent Tapestation or Agilent Femto Pulse. For the initial extractions DNA was diluted, if required, to < 50ng/ ***μ***l and a 1 ***μ***l aliquot run on an Agilent Genomic Tape on a Tapestation instrument according to the manufacturer’s instructions. For the second set of extractions DNA was diluted to 0.25ng/ ***μ***l and a 1 ***μ***l aliquot run on an Agilent Femto Pulse instrument according to the manufacturer’s instructions Electropherograms for each bacterial species can be seen in the supplementary material.

### 5.3. Construction of the synthetic mocks

Two synthetic mocks consisting of all 7 species at 7 different proportions were constructed. For both mocks we targeted 50% A. xylosoxidans, 25% M. morganii, 12% L. richardii, 6% P. aeruginosa, 4% M. wisconsensis, 2% P. vulgaris and 1% S. dysgalactiae based on average Qubit measurements. The first was used for the 1.7kbp, 4.7kbp and 12.8kbp runs and the second for the 10.6kbp runs.

To remove smaller molecules and improve average read lengths a size exclusion step using the Sage Scientific BluePippin was performed. Four 5***μ***g aliquots of the unfragmented mock were run on a High Pass cassette on a BluePippin to remove molecules <15 kbp according to the manufacturer’s instructions, collecting the size selected material in 40***μ***l of running buffer.

To target read N50s around 6kbp a 5***μ***l aliquot of the unfragmented mock in 100 ***μ***l was placed in a G-tube and spun for 2x 1 minute at 10 000 rpm in an Eppendorf 5415 centrifuge. To confirm the size of the fragmented DNA a 1***μ***l aliquot was run on a Agilent Tapestation genomic tape according to the manufacturer’s instructions(see supplementary material).

### 5.4. Library construction and Sequencing

Libraries for the 4.5 kbp, 12.8kbp size exclusion, 10.6 kbp and 16.4 kbp runs were constructed using the Oxford Nanopore Technologies (ONT) SQK-LSK109 kit according to the manufacturer’s instructions except that Kapa beads (Roche, UK) were used to perform the clean up steps rather than Ampure XP beads. To target average sequence reads of 1.7 kbp,100ng of G-tube fragmented mock was used in a 10 ***μ***l reaction using the ONT RAD004 kit according to the manufacturer’s instructions.

In all cases final libraries were sequenced on individual R9.4.1 Rev D 106 flowcells on an ONT GridION.

When targeting successive enrichment of each individual species within the mock, runs were set up with no enrichment for the first hour to ascertain their baseline composition. At the end of the hour the run was stopped and restarted enriching for the next target genome. This process was repeated and sequence data collected for one hour until all seven targets had been selected. For the 1.7kbp, 4.7kbp and 12.4kbp size exclusion runs all 512 pores were chosen to enrich. For the 10.6kbp and 16.4kbp runs pores 1 to 256 were chosen to enrich and pores 257 to 512 were chosen for controls.

Additional runs involved sequencing a 10.6kbp library for 6 hours enriching for S. dysgalactiae only followed by a nuclease flush and re loading the library and running for a further 6 hours enriching for S. dysgalactiae only, running a 10.6kbp library and enriching for M. morganii and collecting for 72 hours, running a 10.6kbp library and enriching for S. dysgalactiae and collecting for 72 hours, running a 10.6kbp library and depleting for M. morganii and collecting for 72 hours and running a 10.6kbp library and depleting for M. morganii, A xylosoxidans and L. richardi and collecting for 72 hours. In each case pores 1 to 256 were chosen to enrich and pores 257 to 512 were chosen for controls.

All sequencing data are available in the European Nucleotide Archive (http://www.ebi.ac.uk/ena) repository under accession number PRJEB44844. ONT run reports, along with a table providing direct links to the ENA runs can be found at https://github.com/richardmleggett/adaptive_sampling.

### 5.5. GridION Adaptive Sampling

For each adaptive sampling run we supply MinKNOW with a reference file containing only the genome of the species we wish to target. This is the reference file that is used to perform the classification of the first ~450bps, upon which the molecule is either sequenced entirely or ejected from the pore. We also use MinKNOW’s “align” function to align all reads to a reference file containing the genomes of all species in the sample. This mapping does not affect the decisions on sequencing or ejecting molecules, and is the mapping we use for our classification. Because the initial classification used to inform the decision on whether to sequence or not must be done very quickly (before the molecule has passed through the pore), it does not necessarily coincide with the more thorough mapping done later. Misclassifications from the initial mapping have a moderate effect on the enrichment we observe.

### 5.6. Adaptive sequencing model web app

A web application was created in R using the “Shiny” library, to allow researchers to see the effect experiment parameters will have on the predicted enrichment, as detailed in Section 2.1. The app is available at https://sr-martin.shinyapps.io/, and the source code can be found in the github repository https://github.com/SR-Martin/Adaptive-Sequencing-Analysis-Scripts.

### 5.7. Bioinformatics analysis

All bespoke scripts used in the analysis were written in Python and are freely available in the github repository https://github.com/SR-Martin/Adaptive-Sequencing-Analysis-Scripts.

Sequences were basecalled during the experiment on the GridION using the MinKNOW software. Mappings of each read to the reference genomes of the seven species in the mock community were also created by MinKNOW. The script analyse_RU.py was used to cross reference the mappings with the reads, and report read and bp statistics for each species, split by channels used for adaptive sampling and all others (when appropriate).

For the analysis of false negatives, the script RU_decision_stats.py was used to parse the adaptive sampling logs created by MinKNOW for each experiment. This script determines the signal sent to the pore for each read and uses these to split the read set into reads that have been ejected from the pore (“unblocked”) and those that were sequenced. These read sets were then cross referenced with the file of mappings and reads were manually binned by species and signal type. The script get_read_stats.py was used to obtain statistics for each read set.

The read length distributions in Figure 1a and the control distributions in Figure 4a were obtained by binning reads by length into bins of size 1,000. For the enrichment distributions in Figure 4a, reads were binned by length into bins of size 100.

In Figure 3.c, yields were normalised by the number of active channels, where active channels were those that sequenced a molecule in the first 30 minutes of the experiment. For the plots of active channels over time (Figure 6b and Figure 7), a channel was defined as active from the beginning of the experiment, up until the time it sequenced its final molecule (as long as it sequenced at least one molecule). Active channels were counted using the script GetActiveChannels.py, with counts each hour for the 72-hour experiments, and every 15 minutes for the 6-hour nuclease flush experiment.

For Figure 6d, the time between two successive target molecules was recorded for each channel using the script GetWaitingTimes.py. For Figure 6c the script GetTimeHist.py was used to get the target yield for channels 1-256 and 257-512 each hour. For the yield plot in Figure 7, a different approach was taken to reduce the effect of the mux scans; the script GetTimeHistFlush.py was used to get the total yield for channels 1-256 and 257-512 in sliding windows every 15 minutes. For the first six 15-minute intervals, the sliding window was the duration of the experiment up to that point. For the remaining intervals, the window was the 90 minutes before. The yield in each window was normalised by its duration. For Figure 5, the script GetYieldByTarget.py was used to determine the yield each hour, split by channels 1-256 and 257-512, and split by reference.

All plots were created in Python using pandas and matplotlib in Jupyter Lab.

### 5.8. MAG assembly

Reads mapping to *S. dysgalactiae* were binned by their start time, with bins containing reads that were sequenced in the first hour, the first two hours, etc. up to all 12 hours, using the script GetReadsByTargetAndTime.py. After 6 hours a nuclease flush was performed. Each bin was assembled with Flye v2.8.1 using the command

~~~
flye --nano-raw <read bin> --genome-size 2.1m
~~~

Assembly statistics were collected with a custom script, and each assembly was compared to the reference genome using dnadiff (part of Mummer v3.23^[31]^).

## Declarations

### Ethics approval and consent to participate

Not applicable

### Consent for publication

Not applicable

### Availability of data and materials

The sequence datasets generated and analysed during the current study are available in the European Nucleotide Archive (http://www.ebi.ac.uk/ena) repository under accession number PRJEB44844.

### Authors’ contributions

RML and MDC designed the study. DH and SH performed the experiments. SM developed the model. SM, YL and RML performed data analysis. RML, MDC, SM, DH and YL wrote the manuscript.

## Acknowledgements

We thank Carl Mayers for early discussions about selective sequencing approaches. This research was supported in part by the NBI Computing Infrastructure for Science Group, which provides technical support and maintenance to Earlham Institute’s high-performance computing cluster and storage systems.

We also thank E Yabuuchi, The Lister Institute, F W Hickamn-Brenner, O Lysenko and F Griffith for depositing the bacterial strains used in this study with the NCTC and making them available to the scientific community.

